# Correcting batch effects in single-cell RNA sequencing data by matching mutual nearest neighbours

**DOI:** 10.1101/165118

**Authors:** Laleh Haghverdi, Aaron T. L. Lun, Michael D. Morgan, John C. Marioni

## Abstract

The presence of batch effects is a well-known problem in experimental data analysis, and single-cell RNA sequencing (scRNA-seq) is no exception. Large-scale scRNA-seq projects that generate data from different laboratories and at different times are rife with batch effects that can fatally compromise integration and interpretation of the data. In such cases, computational batch correction is critical for eliminating uninteresting technical factors and obtaining valid biological conclusions. However, existing methods assume that the composition of cell populations are either known or the same across batches. Here, we present a new strategy for batch correction based on the detection of mutual nearest neighbours in the high-dimensional expression space. Our approach does not rely on pre-defined or equal population compositions across batches, only requiring that a subset of the population be shared between batches. We demonstrate the superiority of our approach over existing methods on a range of simulated and real scRNA-seq data sets. We also show how our method can be applied to integrate scRNA-seq data from two separate studies of early embryonic development.

## 1 Introduction

The decreasing cost of single-cell RNA sequencing experiments (D. A. Jaitin et al. 2014; Klein et al. 2015; Macosko et al. 2015; Gierahn et al. 2017) has encouraged the establishment of large-scale projects such as the Human Cell Atlas, involving the profiling of the transcriptomes of thousands to millions of cells. For such large studies, logistical constraints inevitably dictate that data are generated separately– i.e. at different times and with different operators. Moreover, data may also be generated in multiple laboratories using different cell dissociation and handling protocols, library preparation technologies and/or sequencing platforms. All of these factors result in batch effects (Hicks et al. 2017; Tung et al. 2017), where the expression of cells in one batch differ systematically from those in another batch due to technical reasons. Such differences can mask the underlying biology or introduce spurious structure in the data, and must be corrected prior to further analysis to avoid misleading conclusions.

The most popular existing methods for batch correction are based on linear regression. The limma package provides the removeBatchEffect function (Ritchie et al. 2015), which fits a linear model containing a blocking term for the batch structure to the expression values for each gene. Subsequently, the coefficient for each blocking term is set to zero and the expression values are computed from the remaining terms and residuals, yielding a new expression matrix without batch effects. The ComBat method (Johnson, Li, and Rabinovic 2007) uses a similar strategy but performs an additional step involving empirical Bayes shrinkage of the blocking coefficient estimates. This stabilizes the estimates in the presence of limited replicates by sharing information across genes. Other methods such as RUVseq (Risso et al. 2014) and svaseq (Leek 2014) are also frequently used for batch correction, but focus primarily on identifying unknown factors of variation, e.g., due to unrecorded experimental differences in cell processing. Once these factors are identified, their effects can be regressed out as described previously.

Typical applications of existing batch correction methods to scRNA-seq data assume that the composition of the cell population within each batch is identical. Any systematic differences in the mean gene expression between batches are attributed to technical differences that can be regressed out. However, in practice, population composition is usually not identical across batches in scRNA-seq studies. Even assuming that the same cell types are present in each batch, the abundance of each cell type in the data set can change depending upon subtle differences in cell culture or tissue extraction, dissociation and sorting, etc. Consequently, the estimated coefficients for the batch blocking factors are not purely technical, but contain a non-zero biological component due to differences in composition. Batch correction based on these coefficients will thus yield inaccurate representations of the cellular expression profiles, potentially yielding worse results than if no correction was performed.

Here, we propose a new method for batch correction based on the presence of mutual nearest neighbours (MNNs) between batches. MNNs are identified in the high-dimensional expression space and represent pairs of cells of similar type or state in different batches. The difference in expression values between cells in a MNN pair provides an estimate of the batch effect, which is made more precise by averaging across many such pairs. A correction vector is obtained from the estimated batch effect and applied to the expression values to perform batch correction. Our approach automatically identifies overlaps in population composition between batches and uses only the overlapping subsets for correction, thus avoiding the assumption of equal composition required by other methods. We demonstrate that our approach outperforms existing methods on a range of simulated and real scRNA-seq data sets. We also show how our method can be used to integrate scRNA-seq data sets from two separate studies of mouse gastrulation, demonstrating its utility in a real-life setting.

## 2 Matching mutual nearest neighbours for batch correction

Our approach identifies cells between different experimental batches or replicates that have mutually similar expression profiles. We infer that any differences between these cells in the high-dimensional gene expression space are driven by batch effects and do not represent the underlying biology. Upon correction, multiple batches can be “joined up” into a single data set (Figure 1a).

**Figure 1:**
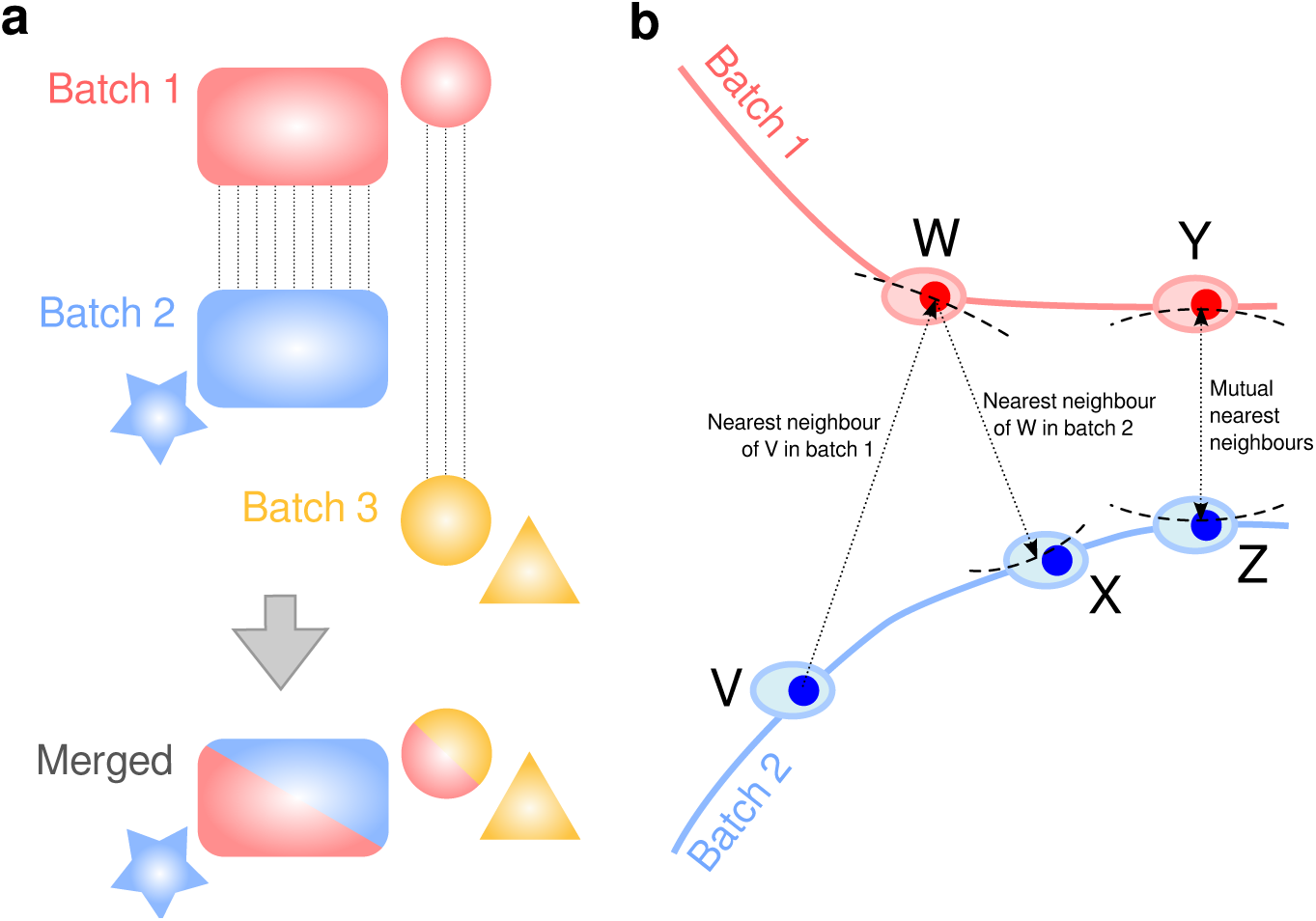
(a) A schematic representing how our MNN method exploits matching subpopulations of cells to learn and apply batch effect correction across several batches. Dotted lines indicate corresponding subpopulations between batches. (b) Identification of matching subpopulations based on mutual nearest neighbours. Cells in each batch are distributed across a smooth manifold, the shape and location of which are specific to each batch (represented by the blue and red lines). For each cell, the set of its nearest neighbours in the other batch are those where the smallest possible radius circle, centered at that cell, lies on the tangent to the other batch manifold (indicated by the dashed arc). Therefore, mutual nearest neighbours are those pairs of cells whose tangent regions are approximately parallel to each other and can only be identified when the two batches are exactly or nearly parallel to each other. To illustrate, for each of two cells W and Y in the red batch, the set of nearest neighbours in the blue batch are identified and marked with a dashed arc. Cell W lies in the set of nearest neighbours of V, but the set of nearest neighbours of W does not include V. This means that W and V are not mutual nearest neighbours. In contrast, cell Y is in the set of nearest neighbours of cell Z and vice versa. Thus, Y and Z are mutual nearest neighbours of each other.

The first step of our method involves global scaling of the data using a cosine normalization. More precisely, if *Y*_*x*_ is the expression vector for cell *x*, we define the cosine normalization as:

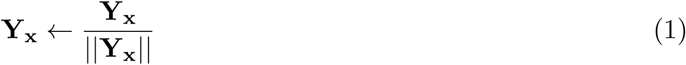

Subsequently, we compute the Euclidean distance between the cosine-normalized expression profiles of pairs of cells (i.e., the cosine distance; see Supplementary Note 1). Cosine distances are appealing as they are scale-independent and robust to technical differences between batches (Bendall et al. 2014). In contrast, Euclidean distances computed using the original expression values are affected by batch-specific factors such as sequencing depth and capture efficiency.

The next step involves identification of **mutual nearest neighbours**. Consider a scRNA-seq experiment consisting of two batches *b* = 1 and 2. For each cell *i*_1_ in batch 1, we find the *k* cells in batch 2 with the smallest distances to *i*_1_, i.e., its *k* nearest neighbours in batch 2. We do the same for each cell in batch 2 to find its *k* nearest neighbours in batch 1. If a pair of cells from each batch are contained in each other’s set of nearest neighbours, those cells are considered to be mutual nearest neighbours (Figure 1b). We interpret these pairs as containing cells that belong to the same cell type or state, despite being generated in different batches. This means that any systematic differences in expression level between cells in MNN pairs must represent the batch effect.

Our use of MNN pairs involves three assumptions: (i) there is at least one cell population that is present in both batches, (ii) the batch effect is orthogonal to the biological subspace, and (iii) batch effects variation is much smaller than the biological effects variation between different cell types (Supplementary Note 2). The biological subspace refers to a set of basis vectors, each of length equal to the number of genes, which represent biological processes. For example, some of these vectors may represent the cell cycle; some vectors may define expression profiles specific to each cell type; while other vectors may represent differentiation or activation states. The true expression profile of each cell can be expressed as a linear sum of these vectors. Meanwhile, the batch effect is represented by a vector of length equal to the number of genes, which is added to the expression profile for each cell in the same batch. Under our assumptions, it is straightforward to show that cells from the same population in different batches will form MNN pairs (Supplementary Note 2). This can be more intuitively understood by realizing that cells from the same population in different batches form parallel hyperplanes with respect to each other (Figure 1b). We also note that the orthogonality assumption is weak for a random one-dimensional batch effect vector in high-dimensional data, especially given that local biological subspaces usually have much lower intrinsic dimensionality than the total number of genes in the data set.

For each MNN pair, a pair-specific batch correction vector is computed as the vector difference between the expression profiles of the paired cells. While a set of biologically relevant genes (e.g. highly variable genes) can facilitate identification of MNNs, the calculation of batch vectors does not need to be performed in the same space. Therefore, we can calculate the batch vectors for a different set of inquiry genes and without the cosine normalisation such that the batch corrected output is on log-transformed scale akin to the input data. A cell-specific batch correction vector is then calculated as a weighted average of these pair-specific vectors, computed using a Gaussian kernel (see Supplementary algorithm box 1). This approach stabilizes the correction for each cell and ensures that it changes smoothly between adjacent cells in the high-dimensional expression space. We emphasize that this correction is performed for all cells, regardless of whether or not they participate in a MNN pair. This means that correction can be performed on all cells in each batch, even if they do not have a corresponding cell type in the other batches.

## 3 Results

### 3.1 MNN correction outperforms existing methods on simulated data

We generated simulated data for a simple scenario with two batches of cells, each consisting of varying proportions of three cell types (Methods 7.1). We applied each batch correction method – our MNN-based correction method, limma and ComBat – to the simulated data, and evaluated the results by inspection of *t*-stochastic nearest-neighbour embedding (*t*-SNE) plots (Maaten and Hinton 2008). Proper removal of the batch effect should result in the formation of three clusters, one for each cell type, where each cluster contains a mixture of cells from both batches. However, we only observed this ideal result after MNN correction (Figure 2). Expression data that was uncorrected or corrected with the other methods exhibited at least one cluster containing cells from only a single batch, indicating that the batch effect was not fully removed. This is fully attributable to the differences in population composition, as discussed earlier. Repeating the simulation with identical proportions of all cell types in each batch yielded equivalent performance for all methods (Supplementary Figure 1).

**Figure 2:**
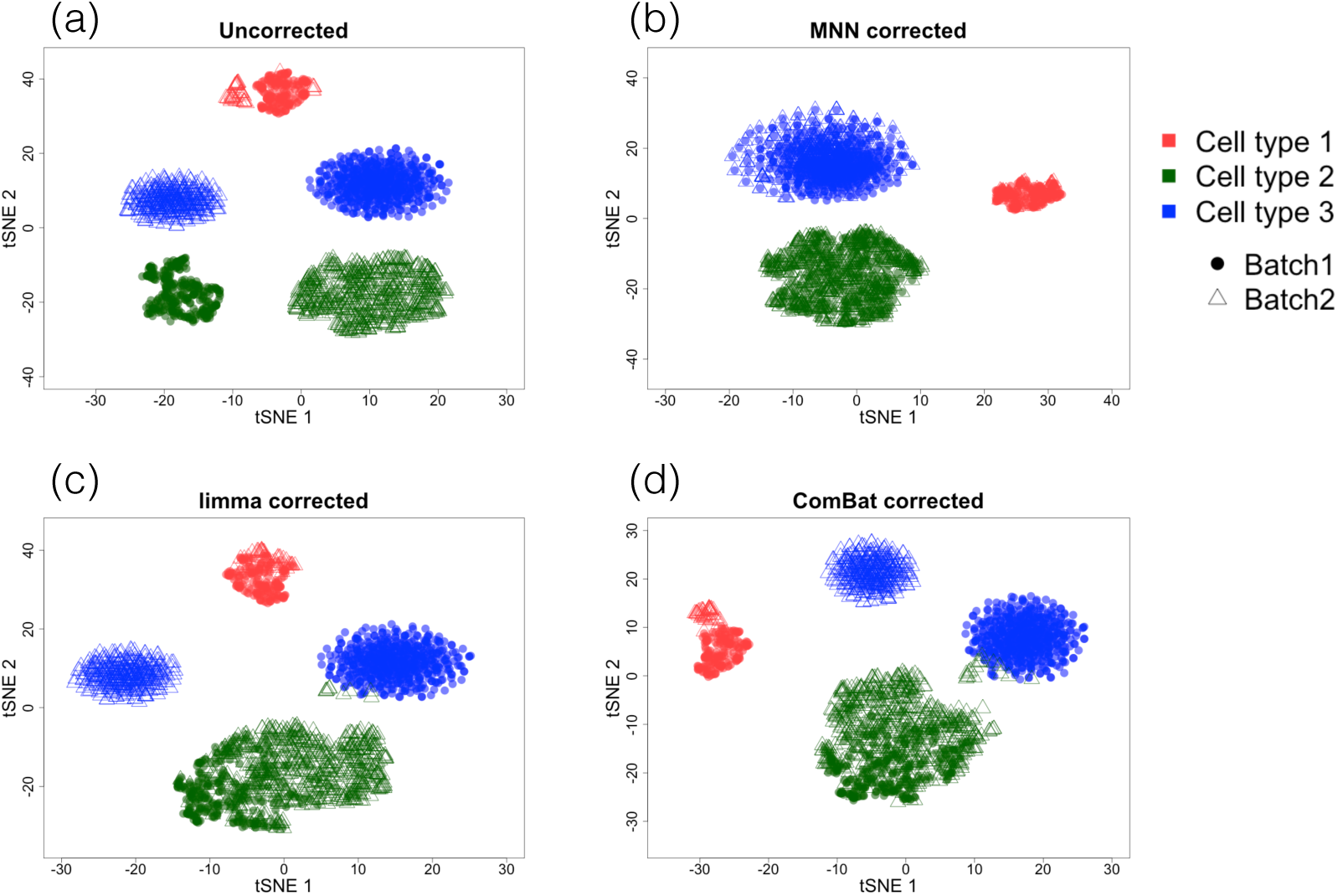
*t*-SNE plots of simulated scRNA-seq data containing two batches of different cell types, (a) before and after correction with (b) our MNN method, (c) limma or (d) ComBat. In this simulation, each batch (closed circle or open triangle) contained different numbers of cells in each of three cell types (specified by colour).

### 3.2 MNN correction outperforms existing methods on haematopoietic data

To demonstrate the applicability of our method on real data, we considered two haematopietic data sets generated in different laboratories using two different scRNA-seq protocols. In the first data set, Nestorowa et al. 2016 used the SMART-seq2 protocol (Picelli et al. 2014) to profile single cells from hematopoietic stem and progenitor cell (HSPC) populations in 12-week-old female mice. Using marker expression profiles from fluorescence-activated cell sorting (FACS), known cell type labels were retrospectively assigned to cells (see Methods Section 7.2). This included multipotent progenitors (MPP), lymphoid-primed multipotent progenitors (LMPP), haematopoietic stem and progenitor cells (HSP), haematopoietic stem cells (HSC), common myeloid progenitors (CMP), granulocyte-monocyte progenitors (GMP), and megakaryocyte-erythrocyte progenitors (MEP). In the second data set, Paul et al. 2015 used the MARS-seq protocol to assess single-cell heterogeneity in myeloid progenitors for 6- to 8-week-old female mice. Again, indexed FACS was used to assign a cell type label (MEP, GMP or CMP) to each cell.

To assess performance, we performed *t*-SNE dimensionality reduction on the (log-scale) highly variable genes of the uncorrected data and the corrected data obtained using each of the three methods (MNN, limma and ComBat) (Figure 3). Only MNN correction was able to correctly merge the cell types that were shared between batches, i.e., CMPs, MEPs and GMPs, while preserving the underlying differentiation hierarchy (Figure 3e, Nestorowa et al. 2016; Paul et al. 2015). In contrast, the shared cell types still clustered by batch after correction with limma or ComBat, indicating that the batch effect had not been completely removed. This is attributable to the differences in cell type composition between batches, consistent with the simulation results. To ensure that these results were not due to an idiosyncrasy of the *t*-SNE method, we repeated our analysis with an alternative dimensionality reduction approach (principal components analysis; PCA) using only the common cell types between the two batches. MNN correction was still the most effective at removing the batch effect compared to the other methods (Supplementary Figure 2).

**Figure 3:**
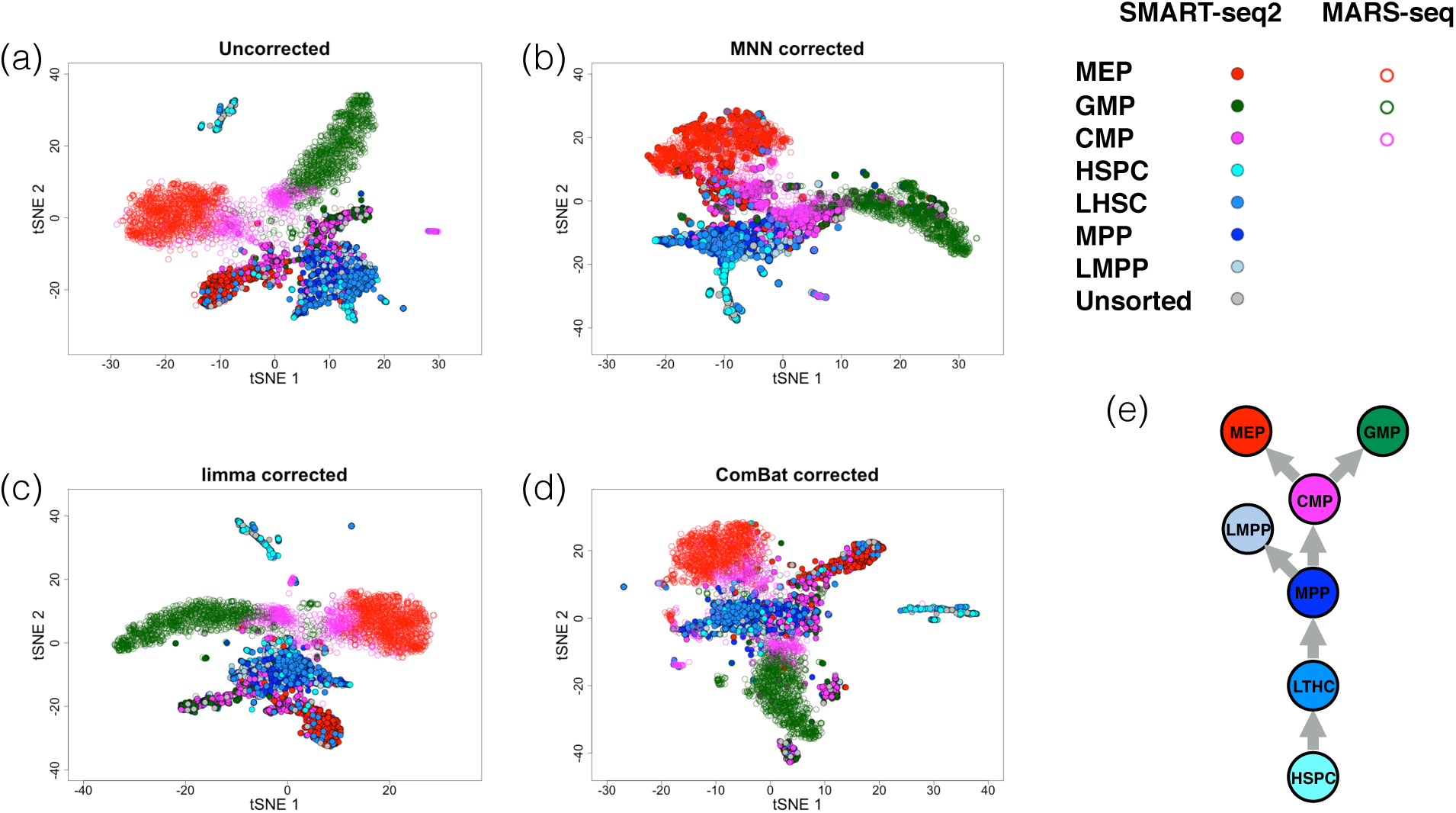
*t*-SNE plots of scRNA-seq count data for cells from the haematopoietic lineage, prepared in two batches using different technologies (SMART-seq2, closed circle; MARS-seq, open circle). Plots were generated (a) before and after batch correction using (b) our MNN method, (c) limma or (d) ComBat. Cells are coloured according to their annotated cell type. (e) The expected hierarchy of haematopoietic cell types.

### 3.3 MNN correction outperforms existing methods on a pancreas data set

We further tested the ability of our method to combine more complex data sets generated using a variety of different methods. Here, we focused on the pancreas as it is a highly heterogeneous tissue with several well-defined cell types. We combined scRNA-seq data on human pancreas cells from four different publicly available datasets (Grün et al. 2016; Muraro et al. 2016; Lawlor et al. 2017; Segerstolpe et al. 2016), generated with two different scRNA-seq protocols (SMART-seq2 and CEL-seq/CEL-seq2). Cell type labels were taken from the provided metadata, or derived by following the methodology described in the original publication – see Methods Section 7.3 for further details of data preprocessing.

We applied MNN, limma and ComBat to the combined data set and examined the corrected data.All three batch correction methods improve the grouping of cells by their cell type labels (Figure 4). This is not surprising, as the discrepancy between cell type composition in the four batches is modest (Supplementary Table 1). However, even a small difference in composition is sufficient to cause ductal and acinar cells to be incorrectly separated following correction with limma or ComBat. By comparison, both cell types are coherently grouped across batches following MNN correction, consistent with the simulation results. To determine the effect of correction on the quality of cell type-based clustering, we assessed cluster separateness by computing the average silhouette widths for each cell type (Fig. 4e, see Methods Section 7.5.1). The average silhouette coefficient after MNN correction is significantly larger than those in the uncorrected, limma- and ComBat-corrected data (pvalue *<* 0.05 Welch’s *t*-test). Thus, MNN correction is able to reduce the between-batch variance within each cell type while preserving differences between cell types. We also computed the entropy of mixing (see Methods Section 7.5.2) to quantify the extent of intermingling of cells from different batches. We observed that batch corrected data using MNN and ComBat show higher entropy of mixing compared to the uncorrected data and batch corrected data using limma (Supplementary Figure 3). This indicates more effective removal of the batch effect for this data set by MNN and ComBat.

**Figure 4:**
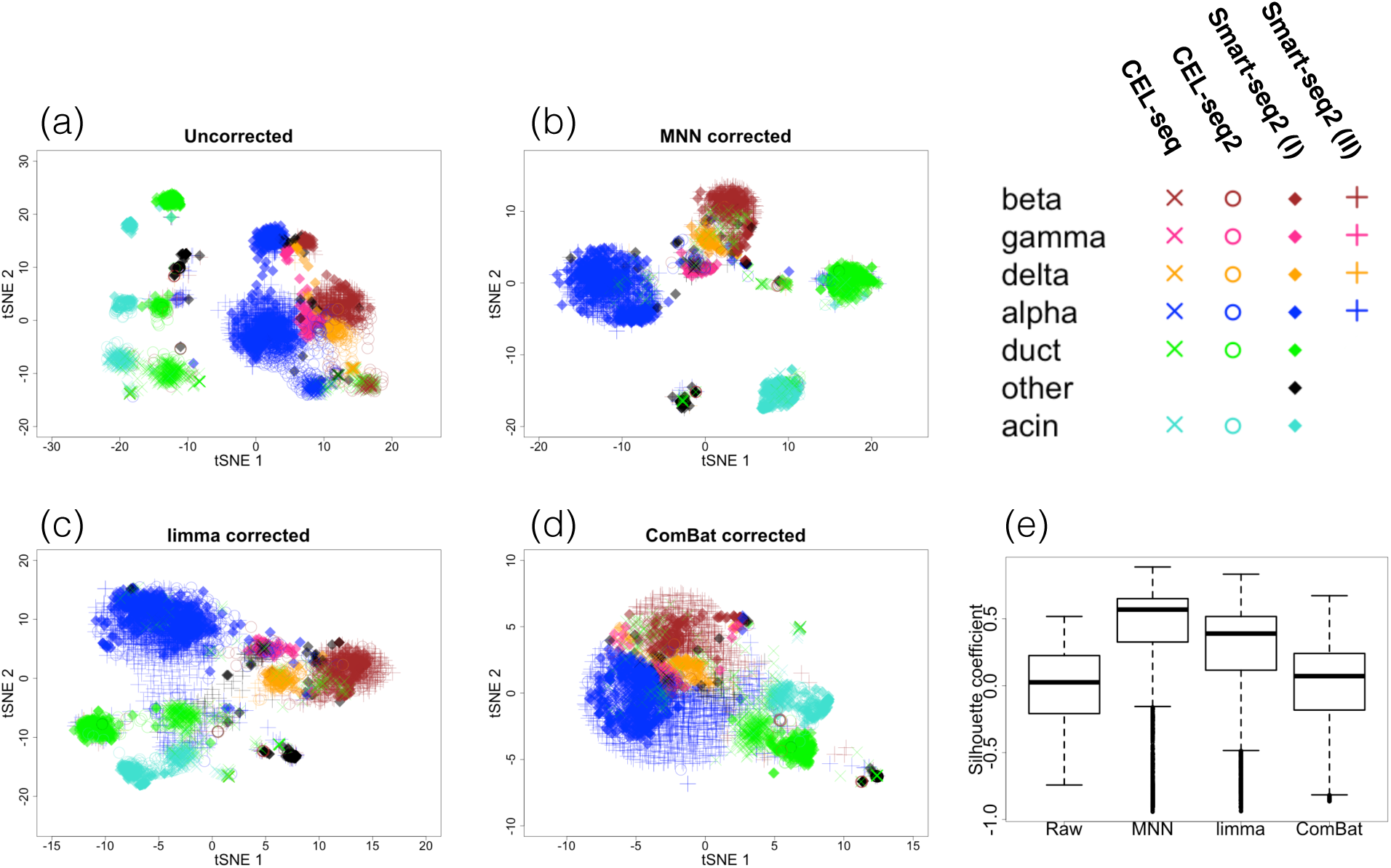
*t*-SNE plots of four batches of pancreas cells assayed using two different platforms, CEL-seq and Smart-seq, for (a) uncorrected (raw) data and the data corrected by (b) our MNN method, (c) limma and (d) ComBat. Different cell types are represented by different colours, while different batches are represented by different point shapes. (e) Boxplots of average silhouette coefficients for separatedness of cell type-based clusters. Whiskers extend to a maximum of *±* 1.5-fold of the interquartile range. Single points denote values outside this range.

#### 3.3.1 Merging data sets with MNN correction improves differential expression analyses

Once batch correction is performed, the corrected expression values can be used in routine downstream analyses such as clustering and differential gene expression identification. To demonstrate, we used the MNN-corrected expression matrix to simultaneously cluster cells from all four pancreas data sets. Our new cluster labels were in broad agreement with the previous cell type assignments based on the individual batches (adjusted Rand index of 0.85). Importantly, we obtained clusters for all batches in a single clustering step. This ensures that the cluster labels are directly comparable between cells in different batches. In contrast, if clustering were performed separately in each batch, there is no guarantee that a (weakly-separated) cluster detected in one batch has a direct counterpart in another batch.

We used our new clusters to perform a differential expression (DE) analysis between the *Δ*-islet cluster and the *γ*-islet cluster. Using cells from all batches, we detected 71 differentially expressed genes at a false discovery rate (FDR) of 5% (Supplementary Figure 4b). This set included the marker genes for the cells included in the analysis (*PPY, SST*), genes involved in pancreatic islet cell development(*PAX6*) and genes recently implicated in *δ*-islet function and type 2 diabetes development (*CD9, HADH*) (Lawlor et al. 2017). For comparison, we repeated the DE analysis using only cells from one batch (Muraro et al. 2016). This yielded only 6 genes at a FDR of 5%, which is a subset of those detected using all cells (Supplementary Figure 4c). Merging data sets is beneficial as it increases the number of cells without extra experimental work; improves statistical power for downstream analyses such as differential gene expression; and in doing so, provides additional biological insights. To this end, our MNN approach is critical as it ensures that merging is performed in a coherent manner.

### 3.4 MNN correction enables better interpretation of early mouse development

Single-cell approaches are particularly amenable to studying early development, where the small number of cells in the developing embryo precludes the use of bulk strategies. One especially important stage of development is gastrulation, when the basic body plan is laid down and many of the cell types that ultimately manifest themselves in the adult are established. Combining data sets generated by different labs at different developmental stages provides a powerful strategy for studying this process.

To demonstrate this, we considered two datasets: one focusing on mouse data from embryonic days E5.5, E6.5 and E6.75 (Mohammed et al. in press), and another focusing on mesodermal diversification by probing cells from E6.5 and E7.0 (Scialdone et al. 2016, Methods Section 7.4). Examination of a *t*-SNE plot generated from the uncorrected data demonstrated that cells clustered by experiment (Fig. 5a), which is especially clear for the E6.5 epiblast cells. After applying our MNN approach, we observed that cells were arranged chronologically from earliest developmental stage (E5.5) to the latest stage (E7.0), consistent with the underlying biology. We overlaid the expression of the key mesodermal regulator Brachyury (encoded by *T*) onto the t-SNE plots. We observed sporadic expression across disparate clusters in the uncorrected data, in contrast to its coherent expression at one end of the chronological progression in the adjusted data. This coherent expression is consistent with the role of Brachyury in early development, where it is up-regulated in cells ingressing through the primitive streak as they commit to a mesodermal cell fate (Scialdone et al. 2016). Our ability to correctly recover this pattern from the combined datasets clearly demonstrates the power of our approach to correct for batch effects and thus increase power to recover true biological signals. Importantly, this coherent expression pattern is also not captured by limma- or ComBat-based corrections (Supplementary Figure 5).

**Figure 5:**
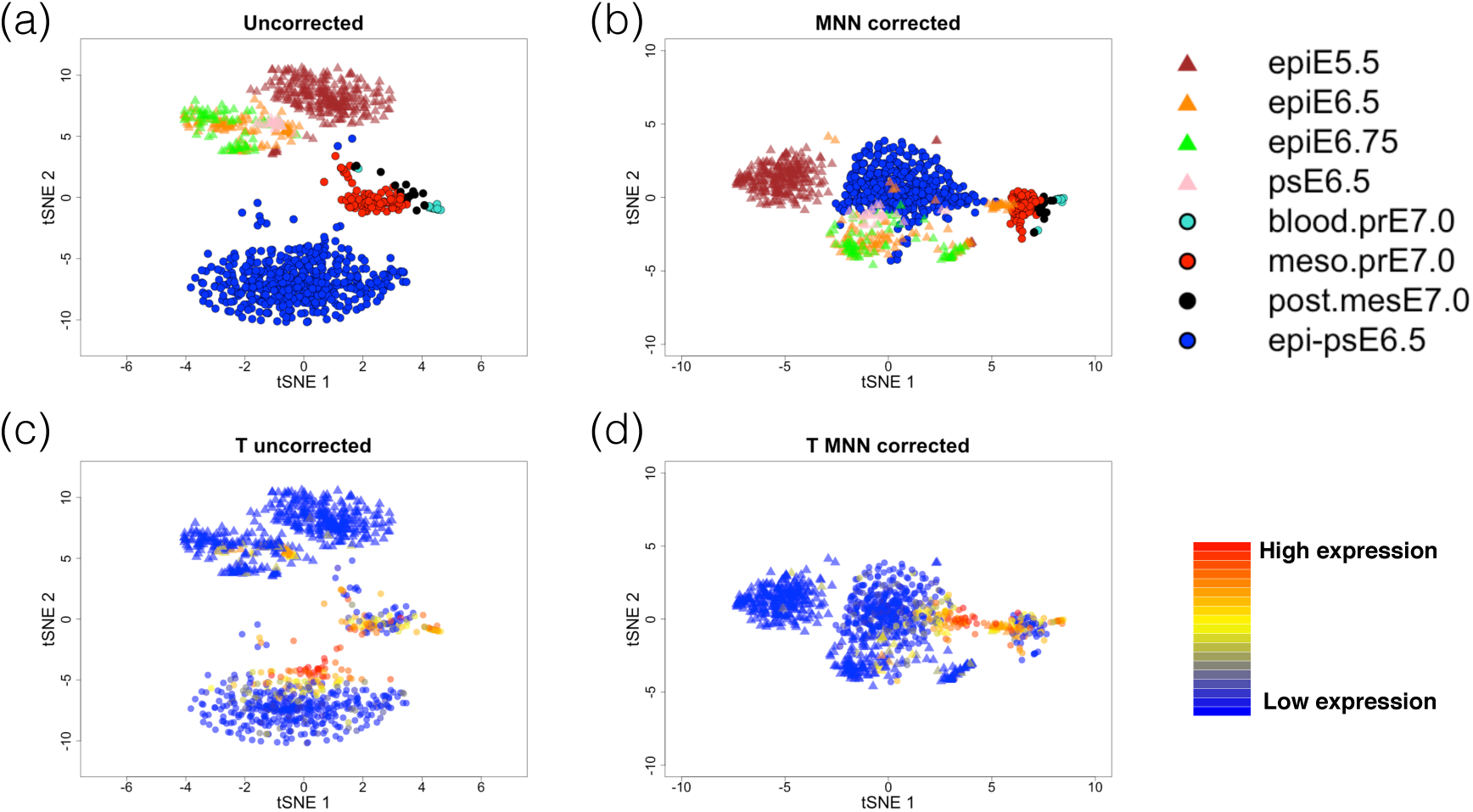
*t*-SNE plots of scRNA-seq data of mouse cells during gastrulation, prepared in two laboratories (Mohammed et al. in press, triangles; Scialdone et al. 2016, circles). Plots were generated (a) before and (a) after MNN correction, with cells coloured by the cell type (epi: epiblast; ps: primitive streak; blood.pr: blood precursor, meso.pr: mesoderm precursor, post.mes: posterior mesoderm, epi-ps: epiblasts transitioning to the primitive streak). Cells were also coloured based on their expression of Brachyury (T), (c) before and (d) after MNN correction.

## 4 Discussion

Proper removal of batch effects is critical for valid data analysis and interpretation of the results. This is especially pertinent as the scale and scope of scRNA-seq experiments increase, exceeding the capacity of data generation within a single batch. To answer the relevant biological questions, merging of data from different batches – generated by different protocols, operators and/or platforms – is required. However, for biological systems that are highly heterogeneous, it is likely that the composition of cell types and states will change across batches, due to stochastic and uncontrollable biological variability.

Existing batch correction methods do not account for differences in cell composition between batches and fail to fully remove the batch effect in such cases. This can lead to misleading conclusions whereby batch-specific clusters are incorrectly interpreted as distinct cell types. We demonstrate that our MNN method is able to successfully remove the batch effect in the presence of differences in composition, using both simulated data and real scRNA-seq data sets. We also show the benefits of merging batches to obtain useful conclusions from a study of gastrulation that was split across several time-based batches.

One prerequisite for our MNN method is that each batch contains at least one shared cell population with another batch. This is necessary for the correct identification of MNN pairs between batches. Batches without any shared structure are inherently difficult to correct, as the batch effects are completely confounded with biological differences. Such cases provide a motivation for using “cell controls”, i.e., an easily reproducible cell population of known composition (say, from a cell line) that is spiked into each sample for the purpose of removing batch effects across samples.

A notable feature of our MNN correction method is that it adjusts for local variations in the batch effects by using a Gaussian kernel. This means that our method can accommodate differences in the size or direction of the batch effect between different cell subpopulations in the high-dimensional space. Such differences are not easily handled by methods based on linear models (as this would require explicit modelling of pre-defined differences between cells, which would defeat the purpose of using scRNA-seq to study population heterogeneity in the first place). This also has some implications for the use of cell controls. Our results for the pancreas data set suggest that considering cell-specific batch effects (the default setting of MNN) rather than a globally constant batch effect for all cells, improves batch removal results (see Supplementary Figure 6). An important consequence is that a single control population might not suffice for accurate estimation of local batch effects. Rather, it may be necessary to use an appropriately mixed population of cells to properly account for local variation.

Batch correction plays a critical role in the sensible interpretation of data from scRNA-seq studies. This includes both small studies, where logistical constraints preclude the generation of data in a single batch; as well as those involving international consortia such as the Human Cell Atlas, where the nature of the project involves scRNA-seq data generation on a variety of related tissues at different times and by multiple laboratories. Our MNN method provides a superior alternative to existing methods for batch correction in the presence of compositional differences between batches. We anticipate that it will improve the rigour of scRNA-seq data analysis and, thus, the quality of the biological conclusions.

## 5 Accession codes

The GEO accession numbers for the published data sets used in this manuscript are provided in the Methods section. An open source software implementation of our MNN method is available as the mnnCorrect function in version 1.4.1 of the scran package on Bioconductor (https://bioconductor.org/packages/scran). All code for producing results and figures in this manuscript are available on Github (<u>https://github.com/MarioniLab/MNN2017</u>).

## 6 Acknowledgement

We are grateful to Fiona K. Hamey (Cambridge Stem Cell Institute) for helpful discussions and to James P. Munro (Center for Mathematical Sciences Cambridge) for helpful mathematical consult. We would also like to thank Maren Büttner (Helmholtz Zentrum München) for helpful discussions.

## 7 Methods

### 7.1 Generation and analysis of simulated data

We considered a three-component Gaussian mixture model in two dimensions (to represent the low dimensional biological subspace), where each mixture component represents a different simulated cell type. Two data sets with *N* = 1000 cells were drawn with different mixing coefficients (0.2*/*0.3*/*0.5 for the first batch and 0.05*/*0.65*/*0.3 for the second batch) for the three cell types. We then projected both data sets to *G* = 100 dimensions using the same random Gaussian matrix, thus simulating high-dimensional gene expression. Batch effects were incorporated by generating a Gaussian random vector for each data set and adding it to the expression profiles for all cells in that data set.

### 7.2 Processing and analysis of the haemaopoetic data sets

Gene expression counts generated by Nestorowa et al. 2016 on the SMART-seq2 platform (1920 cells in total) were downloaded from the NCBI Gene Expression Omnibus (GEO) using the accession number GSE81682. Expression counts generated by Paul et al. 2015 on the MARS-seq platform (10368 cells in total) were obtained from NCBI GEO using the accession GSE72857. For batch correction, we identified a set of 3937 common highly variable genes between the two data sets, by applying the method described by Brennecke et al. 2013 to each data set. For both data sets, we performed library size normalization before log-transforming the normalized expression values. *A priori* cell labels were assigned to each cell based on the original publications.

### 7.3 Processing and analysis of the pancreas data sets

Raw data were obtained from NCBI GEO using the accession numbers GSE81076 (CEL-seq, Grün et al. 2016), GSE85241 (CEL-seq2, Muraro et al. 2016) and GSE86473 (SMART-seq2, Lawlor et al. 2017); or from ArrayExpress, using the accession E-MTAB-5061 (SMART-seq2, Segerstolpe et al. 2016). Count matrices were used as provided by GEO or ArrayExpress, if available. For GSE86473, reads were aligned to the hg38 build of the human genome using STAR version 2.4.2a (Dobin et al. 2013) with default parameters, and assigned to Ensembl build 86 protein-coding genes using featureCounts version 1.4.6 (Liao, Smyth, and Shi 2014).

Quality control was performed on each data set independently to remove poor quality cells (*>*20% of total counts from spike-in transcripts, *<*100,000 reads, *>*40% total counts from ribosomal RNA genes). Sparse cells and genes (*≥* 90% zero values) were also removed, leaving a total of 7236 cells available across all 4 data sets. Normalization of cell-specific biases was performed for each data set using the deconvolution method of Lun, Bach, and Marioni 2016. Counts were divided by size factors to obtain normalised expression values that were log-transformed. Highly variable genes were identified in each data set using the method of Brennecke et al. 2013. We took the intersection of the highly variable genes across all four data sets, resulting in 2496 genes that were used in MNN analysis. The set of 6407 shared gene names among all data sets was used as the inquiry gene set for the batch correction.

Cell type labels for each data set were assigned based on the provided metadata (GSE86473, E-MTAB-5061) or, if the labels were not provided, were inferred from the data using the method employed in the original publication (GSE81076, GSE85241). After correcting for batch effects with each method, we randomly subsampled 1000 cells from each batch prior to applying *t*-SNE, to overcome the *O*(*n*^2^) computational cost involved in calculating all pairwise distances between cells.

To demonstrate the use of the corrected data in downstream analyses, we applied dimensionality reduction (*t*-SNE) to the MNN-corrected expression matrix from the pooled pancreas data sets. We performed *k*-means clustering on the first two *t*-SNE dimensions with the expectation of recovering 8 cell type clusters (see Figure 4 and Supplementary Figure 5). To assign specific cell type labels to these clusters, we examined the expression of the marker genes that were used for cell type assignment in the original publications. Specifically, *GCG* was used to mark *α*-islets, *INS* for *β*-islets, *SST* for *δ*-islets,*PPY* for *γ*-islets, *PRSS1* for acinar cells, *KRT19* for ductal cells, *COL1A1* for mesenchyme cells. Cells in the cluster with the highest expression of each marker gene were assigned to the corresponding cell type. All remaining cells were allocated into an additional “Unassigned/Unknown” cluster.

The differential expression analysis was performed using methods from the limma package (Ritchie et al. 2015). We chose limma as it is fast and valid for large numbers of cells where the central limit theorem is applicable. For the analysis on all cells, we parameterized the design matrix such that each batch-cluster combination formed a separate group in a one-way layout. We used this design to fit a linear model to the normalized log-expression values for each gene, and performed empirical Bayes shrinkage to stabilize the sample variances. A moderated *t*-test was applied to compare the *Δ*- and *γ*-islet clusters across all batches. Specifically, we tested whether the average expression of each cluster across all batches was equal. Differentially expressed genes between the two clusters were defined as those detected at a FDR of 5%. For comparison, we repeated this analysis using only cells from one batch (Muraro et al. 2016). Here, we used a design matrix with a one-way layout constructed from the original cell type assignments. *δ*- and *γ*-islet cell types were directly compared within this batch.

### 7.4 Gastrulation data

Expression counts generated by Mohammed et al. in press on the Hi-seq platform were acquired from the GEO accsession GSE100597. We selected the E5.5, E6.5 and E6.75 stages (466 cells in total) from this data set. Expression counts generated by Scialdone et al. 2016 were downloaded from NCBI GEO using the accession GSE74994. Here, we selected the E6.5 and E7.0 stages (615 cells in total). A set of 13464 matching gene names between the two data sets and the union of highly variable genes according to Brennecke et al. 2013 (3589 genes in total) were used for batch correction. Counts were size-factor normalized using the scran R package and log-transformed prior to downstream analysis.

### 7.5 Quantitative assessment of batch correction methods

#### 7.5.1 Silhouette coefficient

To assess the separatedness of the cell types for the pancreas data, we computed the silhouette coefficient using the kBET package in R. Here, each unique cell type label defines a cluster of cells. Let *〈a*(*i*)*〉* be the average distance of cell *i* to all other cells within the same cluster as *i* and *〈b*(*i*〉) be the average distance of cell *i* to all cells assigned to the neighbouring cluster, i.e., the cluster with the lowest average distance to the cluster of *i*. The silhouette coefficient for cell *i* is defined as:

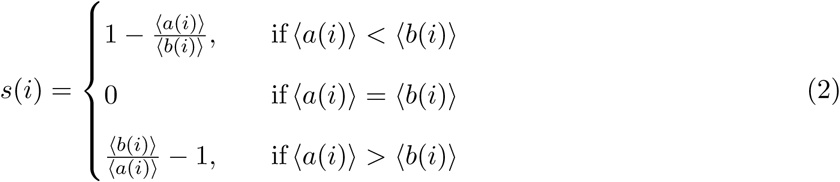

A larger *s*(*i*) implies that the cluster assignment for cell *i* is appropriate, i.e., it is close to other cells in the same cluster yet distant from cells in other clusters. As dimension reduction by *t*-SNE facilitates more reasonable clustering results compared to clustering in the high dimensions, we calculated the silhouette coefficients using distance matrices computed from the *t*-SNE coordinates of each cell in the batch-corrected and the uncorrected data.

#### 7.5.2 Entropy of batch mixing

Entropy of mixing (Brandani et al. 2013) for *c* different batches is defined as:

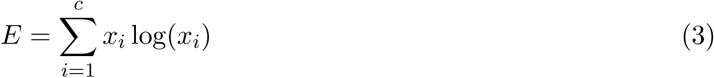

where *x*_*i*_ is the proportion of cells in batch *i* such that 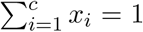We assessed the total entropy of batch mixing on the first two PCs of the batch-corrected and the uncorrected pancreas data sets, using regional mixing entropies according to Equation 3 at the location of 100 randomly chosen cells from all batches. The regional proportion of cells from each batch was defined from the set of 50 nearest neighbours for each randomly chosen cell. The total mixing entropy was then calculated as the sum of the regional entropies. We repeated this for 100 iterations with different randomly chosen cells to generate boxplots of the total entropy (Supplementary Figures 3e and 4f).

